# Theoretical estimates on the expected number of mutations for reconstructing clonal lineage trees

**DOI:** 10.1101/2025.11.20.689642

**Authors:** Nishat Anjum Bristy, Russell Schwartz

## Abstract

Phylogenetics, like many subdisciplines of computational biology, faces a growing challenge of dealing with increasingly large and complicated data sets that have been enabled by ever-improving technologies for sequencing. The issue is particularly acute for studies of somatic evolution, such as of single-cell populations in cancers, where vast single-cell data sets may now identify hundreds of thousands of genetically distinct cells with similar scales of mutations distinguishing them. At the same time, the complexity of the biology of somatic evolution has led to complex phylogeny methods that struggle to scale to even modest data sizes. In this paper, we explore the theoretical and empirical basis for one strategy for managing these large data sets: subsampling mutations to solve the computationally challenging phylogeny problem followed by faster solutions for placement of mutations on a putatively known guide tree. We specifically focus on the fundamental question of determining the number of mutations sufficient to recover the true phylogenetic tree at some level of resolution with high probability. We theoretically analyze variants of several common models that underlie popular tools for building clonal lineage trees. We further evaluate the robustness of these theoretical bounds through simulations of these models, extensions of them, and real biological datasets. The results suggest that modest numbers of mutations suffice to reconstruct clonal trees for typical numbers of clones, supporting the sub-sampling approach as a general strategy for managing the challenges of ever-growing data sets.

## 1 Introduction

Reconstructing evolutionary trees, i.e., phylogenetics, is a fundamental problem in biology aimed at understanding how different entities such as species, genes, or cells diversify from a common ancestor. Applications of phylogenetic trees have branched well beyond their classic use in species/organismal evolution [22,14] to cover applications in single-cell somatic evolution [16] and cell lineage tracing [28]. Phylogenetics has proven a particular powerful tool for studies of somatic evolution in cancer progression [25,20] and the theory and tools developed for that purpose have since helped make possible much broader study of somatic evolution in other diseases [10] or even in putatively healthy tissues [1].

A growing problem in phylogenetic studies, at both species and somatic scales, is scaling to ever-larger data sets. At the species level, the field now needs to tackle databases of tens of thousands to even millions of organisms to be reconciled into single trees [29]. For somatic evolution, single-cell datasets can reach into tens of thousands of single cells per sample [26]. Cancer genomes are particularly problematic because they frequently exhibit phenotypes of hypermutability that can lead to thousands of mutations per tumor distributed across a heterogeneous population of cells [18]. While these vast data sets are crucial for shedding new light on somatic mutation processes they are also algorithmically challenging for phylogenetic inference methods. At the same time, somatic evolution applications have the advantage over typical uses in species trees of a somewhat imprecise definition of taxa (typically species for organismal evolution but genetically similar subpopulations of cells called clones in the somatic case). This leaves the option of solving a problem at a lower resolution (fewer defined clones) if the data do not support finer resolution.

Many methods operate either directly on the reads or on a binary mutation matrix (entities × mutations), where each entry indicates presence or absence of a mutation, which are interpreted in terms of a simplified model of the evolutionary process. Many methods use a “perfect phylogeny” model, which assumes that a mutation is gained once and never lost, equivalent in practice to the “infinite sites” model that there are an infinite number of potentially mutable sites and so each can be mutated only once along the evolutionary history [9]. Under the perfect phylogeny and infinite-sites assumption, the exact phylogeny can be reconstructed in linear time [9]. More complex models based on finite sites assumptions allow a site to be mutated multiple times along the evolutionary history. Models such as Dollo [6] or *k*-Dollo [4] offer a compromise in which we assumes a mutation is gained only once and allowed to be lost at most *k* times along the branches of the phylogeny. Full finite site models may allow arbitrary recurrent mutation or reversion. Particularly as we move to more complicated variant models, such as allowing for complex structural variations (SVs) [3], copy number alterations (CNAs) [21], or combinations of these with more classic single nucleotide variant (SNV) models [7,24], the algorithmic difficulty of the problem may make it infeasible to scale to the sizes of data set now available.

The somatic evolution field has generally dealt with the problem of excessively large data sets heuristically. The most widely used approach is clustering mutations to a small number of equivalence classes that can then be treated as single weighted markers [2]. While valuable, this approach depends on our ability to reliably separate co-occuring mutations by variant allele frequency (VAF), an increasingly difficult proposition for noisy single cell sequencing as mutation numbers grow. Another possibility is the use of consensus tree methods [8], allowing one to apply methods that scale poorly in numbers of markers to subsets of the data and then synthesize the resulting family of trees into a consensus model. In this paper, we focus on a third strategy with precedent in species evolution [5] that we have deployed in our prior work [7]: using a subset of the mutations to solve the harder problem of finding the correct clonal tree topology and then using faster methods to place the remaining mutations onto that putatively known clonal tree. This can be thought of as a variation on a species tree strategy of using a subset of the data to form a reliable backbone tree followed by faster phylogenetic placement methods to place the remainder of the data on the backbone. The mutation sampling version of this problem hinges on the conjecture that if we have a mutation data set that is large compared to the number of clones we need to resolve, then we should be able to find the right tree with a small sample of the data. This paper seeks to put that conjecture on a firmer theoretical basis for somatic evolution models and explore the practical question of how many mutations we need to use in our phylogeny inference to get the right tree topology at a given clonal resolution.

In the remainder of the paper, we assess this conjecture through theoretical and empirical methods. We first determine the bounds for the number of mutations sufficient and necessary for reconstructing evolutionary histories for several variations of phylogeny inference from point mutations (SNVs). For each, we derive theoretical bounds to establish how many mutations one needs to derive the correct tree with confidence. We use experiments on simulated data for these models and more complex variants of them to show that these theoretical bounds yield good results in practice. Our results support the use of the subsampling strategy as a basis for fast and accurate somatic evolution inference in the face of large data sets and costly phylogeny models and provide a basis for further study into extending this strategy of mutation subsampling and placement to more complicated evolutionary and mutation models.

## 2 Preliminaries

We first define some general terminology we will use throughout this paper; other terminology will be introduced as needed. All considered phylogenetic trees 𝒯 are binary trees with node set 𝒱, edge set ℰ, and leaf set *L*. Let *m* = | ℰ|denote the number of edges and *n* = |*L* | the number of leaves. We represent mutation data by a binary matrix *M* ∈ {0, 1} ^*n×s*^, whose rows correspond to entities (e.g., species, observed cells, or clones) and columns correspond to mutations, where *M*_*i,j*_ = 1 indicates that mutation *j* is present in entity *i*. Throughout this section, we treat *M* as generated from an underlying phylogenetic tree *T* according to one of the evolutionary models considered (perfect, 1-Dollo, or coalescent). Let *ℓ*_1_, …, *ℓ*_*m*_ *>* 0 be the edge lengths of 𝒯, and let 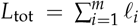 denote the total tree length. Our goal is to determine the expected number of sampled mutations required for every edge of to 𝒯 receive at least one mutation, ensuring that the tree can be reconstructed. Let *p*_*i*_ denote the probability that a mutation lands on edge *i*; given the branch lengths, these probabilities are proportional to the edge lengths,

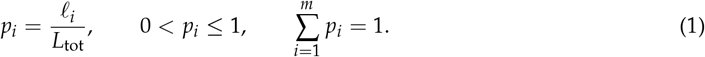

We formalize this as a coverage process over the edges of 𝒯. Each mutation independently selects an edge *i* with probability *p*_*i*_, and we define the “cover time” random variable T as –

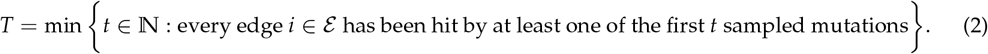

The quantity of interest throughout the paper is the expected cover time 𝔼[*T*], representing the expected number of mutations required for all edges of 𝒯 to be observed at least once.

In the following subsections, we divide the analysis into four parts. We begin with the case of a perfect phylogeny with equal branch lengths and derive the exact number of mutations required for complete coverage, showing that it follows the classic *O*(*m* log *m*) behavior. We then establish general lower and upper bounds for the expected cover time when edge lengths are unequal and sampled from an arbitrary distribution. Finally, we extend the analysis to a loss-supported 1-Dollo phylogeny model and derive the corresponding expected number of sampled mutations.

### Theorem 1

**(Exact cover time for perfect phylogeny with equal branch lengths).** *Let* 𝒯 *be a perfect phylogeny with m edges. Let T be the number of mutations required until every edge has been hit at least once. Then*

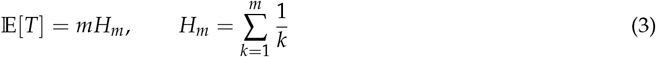

*In particular*, 𝔼[*T*] = *m* log *m* + *γ* + *O*(1) *as m* → ∞, *where γ is the Euler–Mascheroni constant*.

### Proof

Each sampled mutation falls on one of the *m* edges with probability 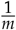. With *p*_*i*_ = 1/*m*, the process of assigning mutations to edges is the classical coupon collector with *m* equiprobable coupons. Let *T*_*i*_ denote the number of additional mutations needed to discover a new edge given that *i* − 1 distinct edges have already been seen (*i* = 0, 1, …, *m*). At that time, the success probability for hitting a previously unseen edge is 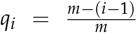. Hence *T*_*i*_ is geometric with mean 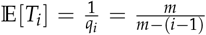. Since 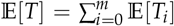 and the *T*_*i*_’s are independent, and thus linearity of expectation suffices.

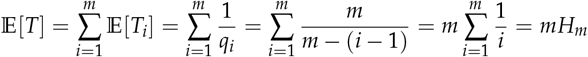

where *H*_*m*_ is the *m* th harmonic number. This can be approximated as 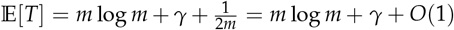, where *γ* ≈ 0.577215 is the Eular-Mashcheroni’s constant. For large *m*, 𝔼[*T*] ≈ *m* log *m* + *γ*. □

We next generalize the problem for unequal edge lengths and thus different probabilities *p*_*i*_ of a sampled mutation getting mapped to an edge. In a classic coupon collectors problem, *p*_*i*_’s are considered fixed parameters. We generalize this to consider *p*_*i*_’s to be random variables following an arbitrary distribution. We will first derive a general upper bound applicable for any such distribution and then consider a special case of the Kingman coalescent with exponentially distributed branch lengths. For the general case, we show in Supplementary Section S1.1 that

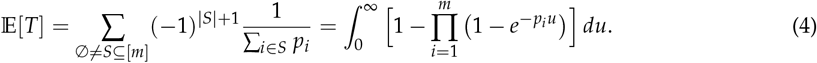

In the equal-probability case *p*_*i*_ = 1/*m*, the integrand simplifies to 1 − (1 − *e*^−*u*/*m*^)^*m*^, and a direct computation gives 𝔼[*T*] = *mH*_*m*_ (Supplementary Lemma S1), where 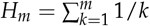 is the *m*th harmonic number. The following theorem generalizes this result for arbitrary edge probabilities.

### Theorem 2

**(Universal upper bound)**. *Let p*_1_, …, *p*_*m*_ *>* 0 *with* ∑_*i*_ *p*_*i*_ = 1, *and* 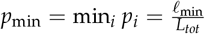. *Then*

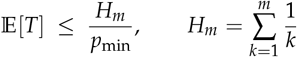

*Equality holds if and only if p*_*i*_ = 1/*m for all edges i*.

### Proof

Starting from Equation (4), note that *x* ↦1 − *e*^−*xu*^ is increasing in *x*, so 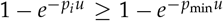 for all *i*. All factors lie in (0, 1], hence

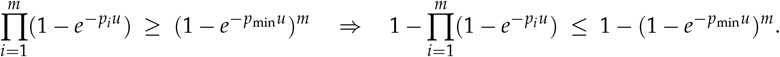

Integrating both sides over *u* ∈ [0, ∞) yeilds and substituting *v* = *p*_min_*u* yields

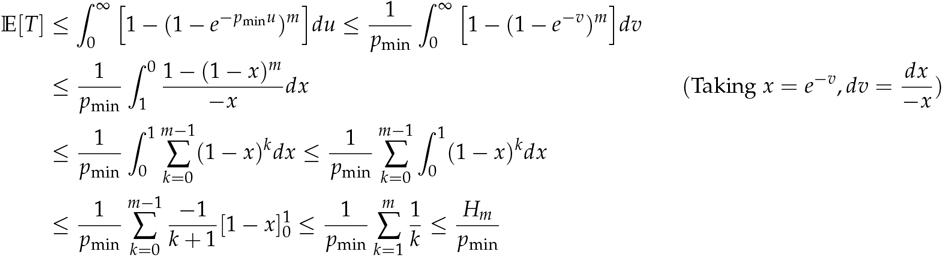

Equality holds only if 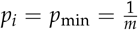 for all *i*. □

### Consequences

If the edge lengths *ℓ*_*i*_ are known, *p*_*i*_ = *ℓ*_*i*_/*L*_tot_ and *ℓ*_min_ = min_*i*_ *ℓ*_*i*_ give the deterministic bound

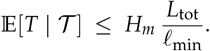

### Special case: Kingman coalescent trees

The Kingman coalescent [11] is a stochastic model of evolution in a neutrally evolving population of constant size. Going backward in time, lineages randomly merge pairwise until a single common ancestor is reached. When *k* lineages are present, the waiting time to the next coalescent event follows an exponential distribution with rate parameter 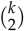. We explore it as an example of an approximation to the edge length ratios we might expect in more realistic clonal trees. We assume 𝒯 to be a random rooted Kingman coalescent binary tree on *n* leaves (*m* = 2*n* − 2 edges) whose edge lengths *ℓ*_*e*_ are independent exponential random variables determined by the aforementioned rates. In this setting, the probabilities *p*_*i*_ = *ℓ*_*i*_/*L*_tot_ are random variables rather than fixed parameters. Taking the expectation over coalescent trees then gives 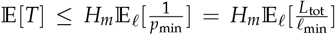. This is the expectation of the ratio of two dependent random variables, which is hard to simplify to a closed-form and often requires knowing the joint distribution (and covariance) of *ℓ*_min_ and *L*_tot_.

To obtain a tractable analytic estimate, we consider a deterministic approximation to the true coalescent in which each coalescent epoch has length fixed to its expected value. In this mean-epoch model, the time between the coalescent events for *k* lineages is 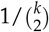. The minimum edge length therefore equals 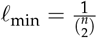 and the total expected tree length is *L*_tot_ = 2*H*_*n*_ − _1_, as we prove in Supplementary Lemma S2. Applying Theorem 2 with *p*_min_ = *ℓ*_min_/*L*_tot_ yields

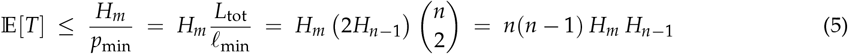

Since *H*_*n*_ = Θ(log *n*), this leads to an asymptotic upper bound of *O n*^2^(log *n*)^2^ . Thus, under this deterministic mean-epoch approximation, the expected number of mutations required to cover all branches of a coalescent tree scales quadratically with the number of lineages, up to logarithmic factors.

Next, we extend the analysis to evolutionary models that permit mutation losses. In cancer evolution, gained mutations are frequently lost due to independent events or through larger-scale copy number deletions [12]. To account for this, the *K*-Dollo model assumes that each mutation may be gained once but can be lost up to *K* times along the phylogeny. The perfect phylogeny corresponds to the special case *K* = 0. We derive the exact expected number of sampled mutations required for full edge coverage under the *1-Dollo model* (*K* = 1), leaving the general *K*-Dollo case as a direction for future work.

#### Theorem 3

**(Expected cover time under the 1-Dollo model)**. *Under the 1-Dollo assumption, each mutation is gained once and may be lost at most once. A single mutation can therefore cover either one previously uncovered edge (gain-only) or two uncovered edges (gain–loss). Let p* = *P*_loss_ ∈ [0, 1) *denote the probability that a gain undergoes loss. Then the expected number of mutations required to cover all m edges is*

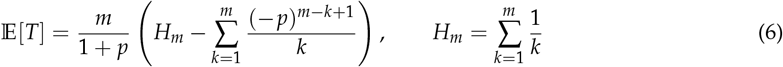

#### Proof

Let’s consider the process of sequentially assigning mutations to the branches of . 𝒯 Let *i* denote the number of edges that are already covered by at least one mutation (equivalently, the number of mutations mapped so far). When a new mutation is sampled, three outcomes are possible: it hits (1) an uncovered gain-only edge, covering one new edge; (2) an uncovered gain–loss pair, covering two new edges; or an already covered edge, covering none. The corresponding state transitions are shown in Figure 1.

**Fig. 1:**
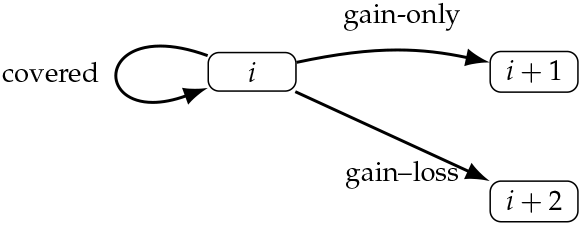
State transitions under the 1-Dollo model. Each new mutation may cover one (gain-only) or two (gain–loss) new edges, or none (already covered).

Let 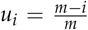 be the probability that the sampled mutation lands on an uncovered edge. Then, *u*_*i*_ *p* will be the probability that the mapped mutation will also have a loss associated with it. Let *P*_1_ = *u*_*i*_(1 − *p*), *P*_2_ = *u*_*i*_ *p, P*_0_ = 1 − *u*_*i*_, corresponding respectively to transitions *i* → *i* + 1, *i* → *i* + 2, and *i* → *i*. Let *E*_*i*_ = 𝔼[*T*_*i*_] denote the expected number of remaining mutations given *i* covered edges. This follows the recurrence

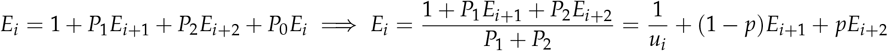

where, *E*_*m*_ = 0 is the base case. Let *r* = *m* − *i* be the number of uncovered edges and define *A*_*r*_ = *E*_*m*−*r*_, where *A*_*r*_ is a reparameterization of *E*_*m*−*r*_ denoting the expected number of mutations when *r* edges are remaining to be mapped. Then, *u*_*i*_ = *r*/*m* and 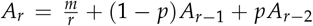 where the base cases are *A*_0_ = 0, *A*_1_ = *m*. Defining *S*_*r*_ = *A*_*r*_ − *A*^*r*−1^ gives 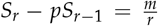 with base case, *S*_1_ = *m*. Solving the recurrence, we get:

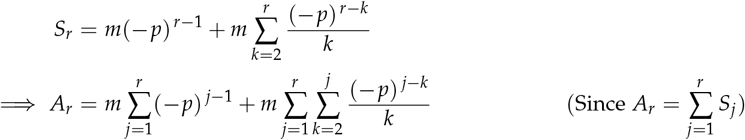

The first term is geometric, so 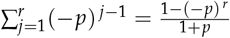 . For the double sum,

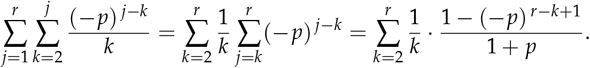

Hence

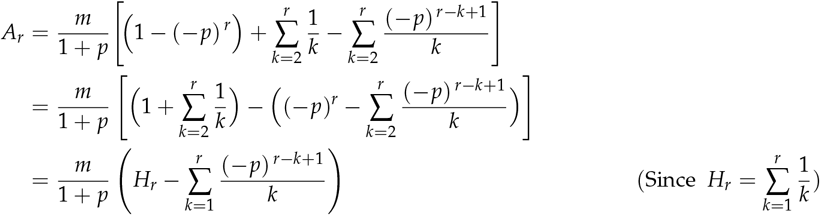

Starting from no covered edges (*i* = 0, *r* = *m*),□

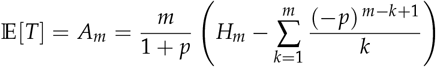

#### Corollary 1

**(No-loss case, i.e. perfect phylogeny)**. *For p* = 0 *(mutations gained once and never lost), the inner sum vanishes and*

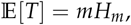

*recovering the classical coupon-collector expectation*.

### Code Availability

All scripts used for simulation are implemented in Python 3 and available at https://github.com/CMUSchwartzLab/mutation-subsampling.git

## 3 Experimental Results

We have conducted experiments on simulated and real variation data designed to validate the correctness of the bounds shown in this paper, assess how pessimistic they are in practice, and test how well they extend to more complex but realistic evolutionary scenarios beyond those covered in the proofs. We first validated the robustness of the bounds using simulated data and assessed tree reconstruction accuracies with respect to SPhyR [4] and PAUP [27] for perfect and 1-Dollo phylogenies, and with PAUP for coalescent-derived phylogenies. We further used simulated data to assess the k-Dollo case and the full coalescent model empirically. Additionally, we evaluated the effectiveness of these bounds on real cancer patient samples. For these cases, we limit the comparisons to SNVs as in our theoretical bounds, although the true data contains multiple mutation types, including SNVs, CNAs, and SVs.

### Evaluation criteria

To assess the quality of the estimated tree topologies on simulated datasets, we compared them with ground-truth trees using *normalized Robinson-Foulds (RF) distance* [19]. To evaluate branch-length accuracy, especially under mutation subsampling, we further computed the *mean absolute error (MAE)* between the true versus estimated branch lengths after normalizing the branch lengths for each tree by that tree’s total branch length, computed over all pairs of leaves in the *n*-leaf trees. Additionally, to evaluate the accuracy of mutation placement on the inferred trees, we compute the *pairwise ancestral relationship accuracy* as described in Supplementary Section S2.1.

### 3.1 Evaluation on simulated data

To evaluate how much phylogeny quality is affected by mutation counts across the range from the lower and upper bounds established theoretically, we simulated single-cell WGS data with SNVs based on known ground-truth tumor and coalescent phylogenies. We explain the simulation process for perfect, *K*-Dollo and coalescent phylogenies in detail in the following sections.

#### Perfect and 1-dollo phylogeny simulations

To evaluate the robustness of our bounds for perfect and 1-Dollo simulations, we first generated random full binary perfect phylogenies with *n* ∈ { 2, 3, 4, 5, 6, 10, 15, 20} leaves. This corresponds to (2*n* − 1) ∈ {3, 5, 7, 9, 11, 19, 29, 39 } total nodes or clones in the tree. Since a full binary tree with *n* leaves has *m* = 2*n* − 2 edges, we generated *mH*_*m*_ mutations and mapped each of these *mH*_*m*_ mutations to a uniformly chosen edge for each full binary perfect phylogeny, 𝒯^∗^. After assigning all mutations, we counted how many branches of the perfect phylogeny remained without any mutation. We simulated 10 replicates for each value of *n*. Figure S1 shows the distribution of the number of unmapped branches. The results indicate that using the perfect-phylogeny bound almost always results in every branch receiving at least one mutation. We ran SPhyR [4] on the generated perfect phylogenies with *mH*_*m*_ mutations. Figure 2 shows the results of SPhyR for perfect phylogenies with the aforementioned number of clones. In each case, we reconstruct the ground truth tree with SPhyR, resulting in zero normalized RF distance and pairwise ancestral accuracies = 1.

**Fig. 2:**
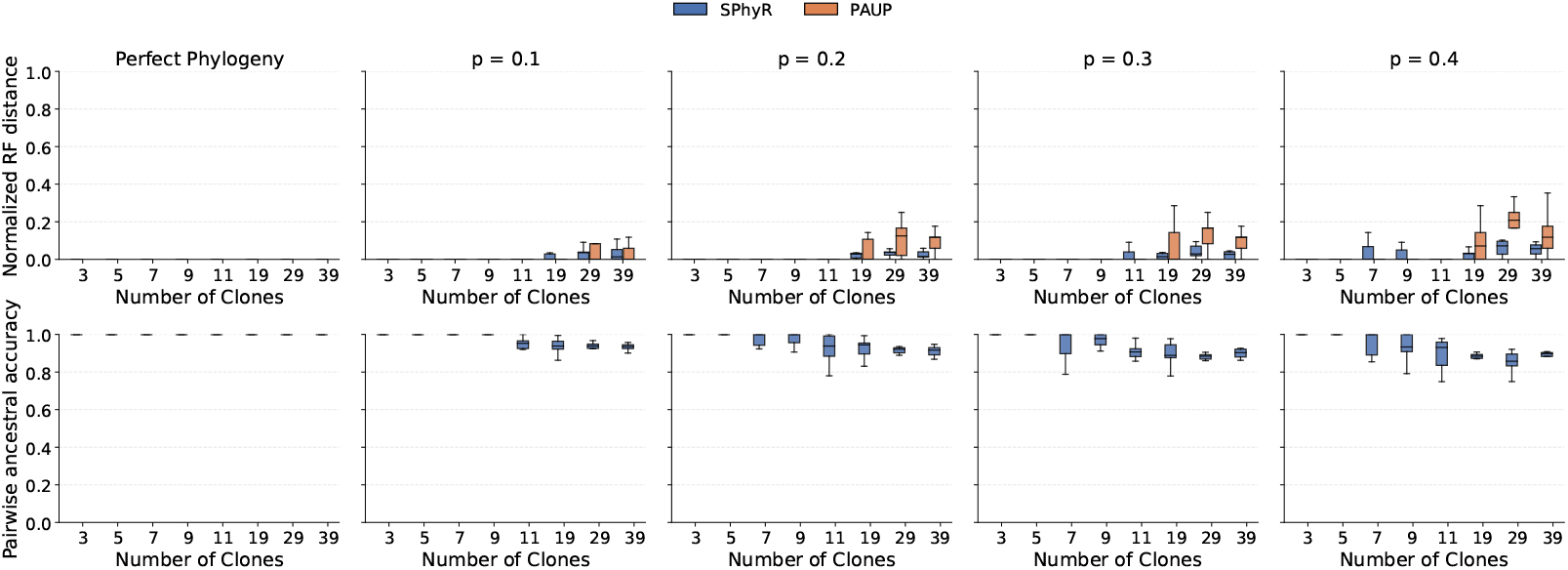
Results for perfect and 1-Dollo phylogeny reconstruction with SPhyR, where mutations are sampled using Equations 3 and 6 respectively for different number of clones *N* ∈ {3, 5, 7, 9, 11, 19, 29, 39}. For the 1-Dollo phylogenies, we consider different loss probabilities, *p* ∈ {0.1, 0.2, 0.3, 0.4}.

To generate 1-Dollo phylogenies, we extended each perfect phylogeny 𝒯^∗^ with *mH*_*m*_ mutations by introducing single-loss events. For each mutation, we independently introduced a loss with probability *p*, removing the mutation and all of its descendants. This produces 1-Dollo instances that contain both gains and single independent losses while preserving the structure of 𝒯^∗^. We then sample a subset of mutations according to Equation (6), run SPhyR [4] and PAUP [27] on the sampled data, and compare the reconstructed trees against the full 1-Dollo phylogeny. Figure 2 shows the 1-Dollo phylogeny reconstruction accuracies. We run 10 replicates for each loss probability *p* ∈ {0.1, 0.2, 0.3, 0.4} and vary the number of clones as {3, 5, 7, 9, 11, 19, 29, 39} . For each simulated instance, we run SPhyR [4] and compare the reconstructed trees using normalized Robinson–Foulds (RF) distances (top panel) and pairwise ancestral relationship accuracies (bottom panel). As the number of clones increases, the normalized RF distance increases slightly, although the reconstructed trees remain highly accurate. We see similar results when evaluating mutation placements via pairwise ancestral relationship accuracy. With three and five clones, there is little difference between the sampled and full 1-Dollo trees, supporting the accuracy of the reconstructed trees for both cases. However, as the number of clones increases, the gap between the mutations in the full and sampled 1-Dollo trees widens, increasing the complexity for SPhyR and leading to larger reconstruction errors. We see similar results for different values of the mutation loss probability, *p* ∈ {0.1, 0.2, 0.3, 0.4} . According to Equation (6), higher loss probabilities result in fewer sampled mutations, which explains the gradual increase in RF distances and the corresponding decrease in pairwise ancestral accuracies as *p* increases. Nonetheless, both topology and phylogenetic placement remain highly accurate on the ranges of clones considered, well beyond the 10 clones typically resolved in tumor phylogeny studies.

#### Empirical evaluation on k-Dollo phylogenies

We generalize beyond the theoretical bounds of the 1-Dollo phylogenies to a more realistic scenario for somatic evolutionary trees, simulating *K*-Dollo phylogenies following SPhyR [4] and the similar method as described in the previous section, by introducing losses in the perfect phylogeny, 𝒯^∗^ for each mutation with loss probability *p*, allowing at most *k* losses per mutation. Specifically, we iteratively select a mutation and introduce a loss with probability *p* if the mutation currently has at most (*k* − 1) losses. Once a loss is introduced for mutation *i*, all descendants of the corresponding node are set to carry that loss.

Since we do not derive an analytical bound for *k*-Dollo phylogenies, we instead provide empirical results for *k* ∈ {1, 2, 3, 4 }. For each *k*, we vary the number of sampled mutations as a multiple *x* of *mH*_*m*_ ( *x* × *mH*_*m*_), where *x* ∈ {0.05, 0.1, 0.5, 0.7, 1, 2, 3} and run SPhyR on the simualted instances with its *K*-Dollo setting. Figure 3 summarizes the empirical estimates. Across all *k* values, we observe that the normalized RF distances decrease and the pairwise ancestral relationship accuracies increase beginning around *x* = 0.7 - 1.0, indicating that the trees become reliably reconstructable once the number of sampled mutations reaches this range for a loss probability of *p* = 0.1. When we calculated the theoretical ratio for x, we derived the values {1.0, 0.89, 0.93, 0.91, 0.9, 0.92, 0.92} respectively for clones ∈ {3, 5, 7, 9, 11, 19, 29}, which matches the empirical results from the *k* = 1 case. We additionally simulated *k*-Dollo phylogenies with higher loss probabilities *p* ∈ { 0.2, 0.3, 0.4} . In the supplementary figures (Figures S2, S3, and S4), we observe similar trends to the *p* = 0.1 case. However, as both the loss probability and *k* increase, SPhyR requires larger number of sampled mutations to infer accurate phylogenies.

**Fig. 3:**
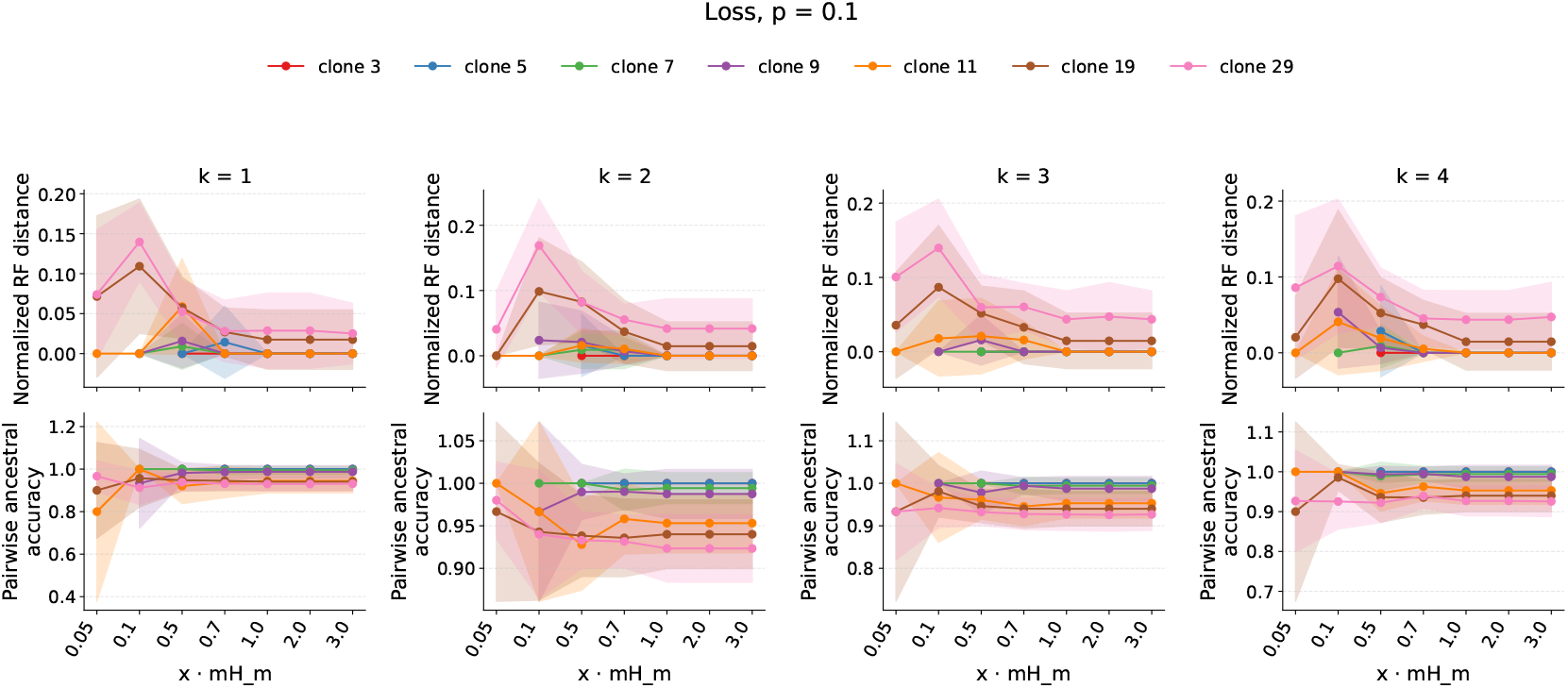
Results for k-Dollo phylogeny reconstruction with SPhyR [4]. The figure shows normalized phylogeny inference (top) and pairwise ancestral accuracy (bottom) across a range of *k*, clone numbers, and sampling rates for loss probability, *p* = 0.1.

#### Coalescent tree simulations

To simulate coalescent trees with *n* leaves representing *n* observed clones, we use CellCoal [17], which starts by simulating a genealogy for the sampled cells under the neutral coalescent model with an exponential population growth model [23] for a defined parameterization of the coalescent for cancer cell samples [15]. A healthy somatic cell is considered to define the “outgroup branch” for the cell phylogeny. We simulate 10 replicates for each of *n* ∈ {3, 5, 7, 9, 11} clone trees, running CellCoal with 0/1 alphabet under the infinite sites model (ISM).

We considered two settings of CellCoal. (1) In the first setting, we explicitly defined the coalescent parameters, setting the effective population size to 10^5^, the exponential growth rate to 10^−5^ and 10,000 variant sites, and generated trees whose branch lengths were exponential random variables following the Kingman Coalescent model. (2) In the second setting, we generated trees with branch lengths fixed to the expected mean-epoch lengths under the coalescent model, such that each inter-coalescent interval corresponded exactly to its theoretical mean. For both settings, the simulated healthy outgroup cell is generated from a reference genome and includes representative germline SNP positions, which we exclude from downstream analyses to ensure only somatic variants are considered.

After the simulation, we run PAUP [27] to reconstruct phylogenies from the coalescent-simulated data. To test how phylogeny reconstruction performs with different numbers of sampled mutations, we first run PAUP with all the non-SNP positions from the coalescent simulations. We then repeat the analysis with *n*(*n* −1)*H*_*m*_ *H*_*n* − 1_ subsampled mutations, following the mean-epoch coalescent bound in Equation (5).

Figures 4(a)–(b) show the normalized Robinson–Foulds (RF) distance and the mean absolute error (MAE) of normalized branch lengths for the random coalescent-branch simulations, while Figures 4(c)–(d) show the corresponding results for trees with deterministic mean-epoch branch lengths. In both cases, black lines represent tree reconstructions using the full set of mutations, whereas red lines correspond to subsampling exactly *n*(*n*−1)*H*_*m*_ *H*_*n*−1_ mutations. We see that across both simulation models, tree topologies were reconstructed with high accuracies even under mutation subsampling, indicating that the predicted number of mutations is sufficient to recover the correct tree structures. However, the branch-length estimation was more challenging for trees with random coalescent branch lengths, PAUP exhibited higher variability and error, whereas trees with deterministic mean-epoch lengths showed substantially more consistent and accurate branch-length estimation.

**Fig. 4:**
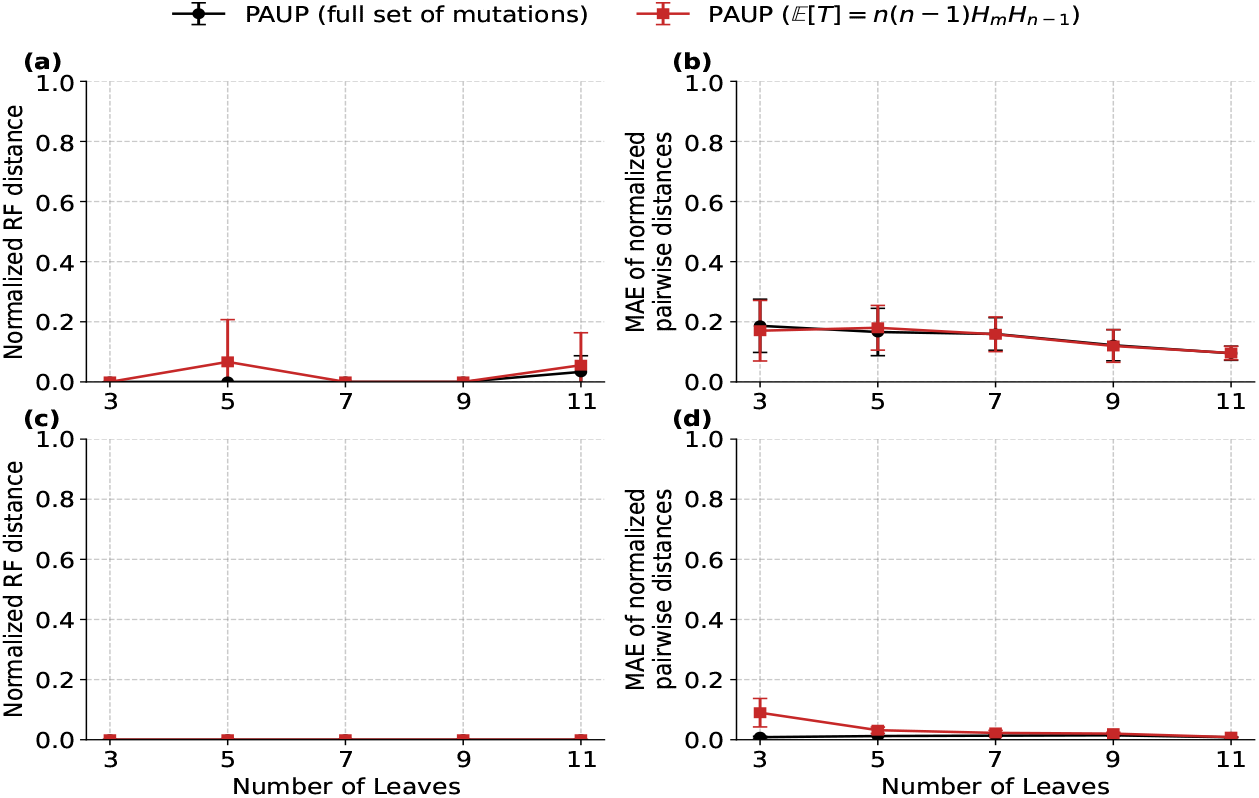
Results of coalescent phylogeny reconstruction using the full set (black lines) and subsampled set (red lines) of mutations across varying numbers of leaves, *n* ∈ 3, 5, 7, 9, 11. (a) and (b) show the normalized Robinson–Foulds (RF) distance and the mean absolute error (MAE) in branch-length estimation for random coalescent simulations following the Kingman model. (c) and (d) show the corresponding RF distance and MAE for coalescent simulations with branch lengths fixed to the expected mean-epoch lengths under the coalescent model.

### 3.2 Evaluation on a metastatic colorectal cancer patient

To evaluate the robustness of our theoretical bounds on a real cancer dataset, we analyzed the metastatic colorectal cancer dataset (CRC1) from Leung et al. [13], which includes 178 single cells and 16 single-nucleotide variants (SNVs) identified through targeted sequencing of 1000 cancer-associated genes. Since the theoretical bounds exceeded the number of SNVs available for reconstructing a perfect phylogeny with more than four clusters, we tested 10 replicates of three and four taxa trees, comparing reconstructions from the sampled set of mutations versus the full set of mutations. We additionally restricted the number of mutations to the maximum available in the dataset and reconstructed five and six clusters trees using PAUP [27]. In each case, PAUP produced identical trees from the sampled versus full mutation sets, resulting in a normalized Robinson–Foulds (RF) distance of 0. To further assess similarity, we compared the L1 distance of the pairwise branch lengths normalized by total tree length for the full and sampled clonal trees. As shown in Figure 5, the number of sampled mutations increases with the number of taxa, while the L1 distance between the branch-length ratios of the full and sampled trees decreases, indicating increasing structural similarity. The results confirm that following the theoretical bounds reliably yields equivalent topologies to the full data set, although with some error in inferred edge lengths.

**Fig. 5:**
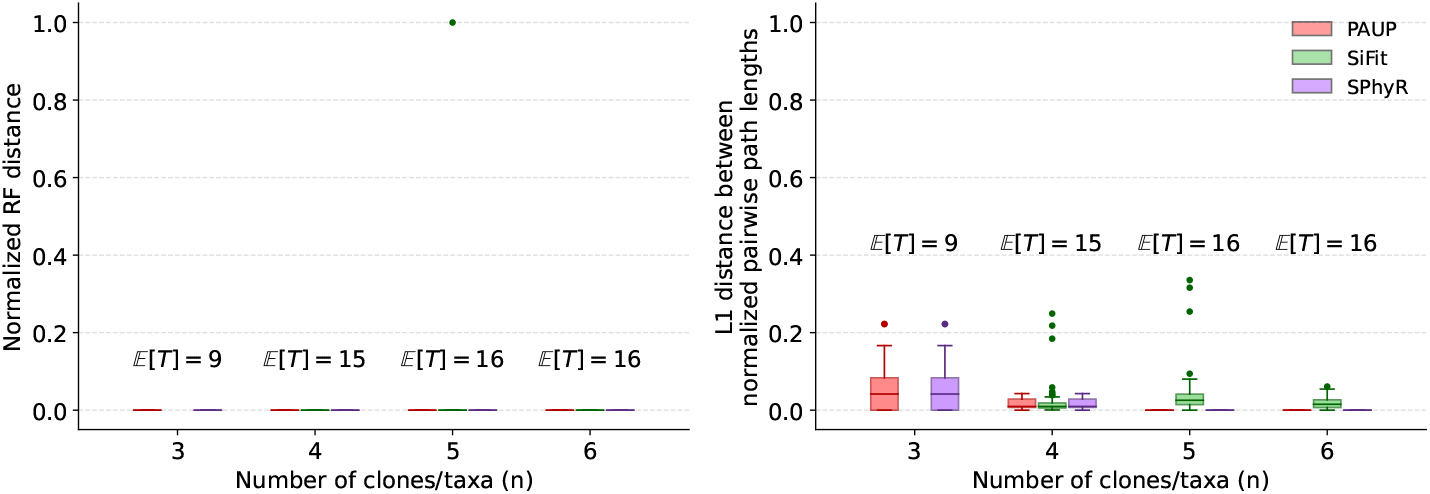
Results on the CRC1 patient dataset [13] for *n* ∈ 3, 4, 5, 6 taxa (clusters). We ran PAUP [27] using the neighbor-joining method for 10 replicates of the sampled mutations with 𝔼 [*T*] ∈ 9, 15, 16, 16 for *n* ∈ 3, 4, 5, 6, respectively.

## 4 Discussion

This paper examined the problem of how many mutations one needs to reliably reconstruct clonal lineage trees from SNV data under various evolutionary models. The question is motivated by a practical problem of how to solve for complex and computationally intensive evolutionary models needed in somatic clone tree applications in the face of increasingly large variation data sets. We established theoretical bounds for some simplified evolutionary models commonly used in clonal lineage studies, showing that modest numbers of variants are sufficient to derive the correct tree topologies for realistic numbers of clones. Empirical studies with simulated and real tumor data confirm that these bounds provide good guidance in practice despite the simplified evolutionary models from which they derive.

This work provides a firmer basis for a practical strategy of dealing with large variant sets and algorithms that scale poorly in their size by subsampling to smaller numbers of variants to derive the correct tree followed by using faster algorithms to place the remaining variants on the tree. Considerable room remains, though, to extend the work further. The theory so far covers only some relatively simplified evolutionary models typical of those assumed in somatic phylogeny methods, but might be extended to more involved models that better capture the true biology. Results on simulated and real data suggest that the bounds we establish hold at least approximately for more realistic data. The present work has only looked at SNV data, and both the theoretical and empirical studies ought to be extended to cover the harder cases of evolution by CNA and SV events important in somatic evolution processes. There is likely also room for improvement in algorithms for both inference of topologies from small variant sets and on fast placement of unsampled variants on that topology, as well as other extensions to more effectively use unsampled variants to validate topologies or assess uncertainty within them. Furthermore, there may be other uses of the derived bounds to explore. For example, while we focused on the scenario of how to subsample mutations when they are available in excess, there is a parallel problem of how to define the maximum level of resolution of clonal structure our data can support when mutations are sparse.

