## Supplementary Materials for "Theoretical estimates on the expected number of mutations for reconstructing clonal lineage trees"

### S1 Supplementary Methods

#### S1.1 Derivation of the Integral Identity for $\mathbb{E}[T]$

**Theorem S1 (Integral Identity for Expected Cover Time under Nonuniform Branch Probabilities).** *Let $T$ denote the number of independent mutation events required until every edge of a phylogenetic tree with $m$ edges has received at least one mutation. Assuming that each mutation independently targets edge $i$ with probability $p_i > 0$, where $\sum_{i=1}^{m} p_i = 1$. Then the expected cover time $\mathbb{E}[T]$ admits the integral representation*

$$\mathbb{E}[T] = \int_0^\infty \left[1 - \prod_{i=1}^{m} \left(1 - e^{-p_i u}\right)\right] du \tag{S1}$$

*Proof.* We sketch the derivation, following a generalized coupon–collector like argument for nonuniform sampling probabilities. For any nonnegative integer-valued random variable $T$, let $\Pr(T > t)$ is the probability that after $t$ mutations at least one edge remains unmapped. Thus we have:

$$\mathbb{E}[T] = \sum_{t \geq 0} \Pr(T > t) \tag{S2}$$

Let $A_i(t)$ denote the event that edge $i$ has not been hit by any of the first $t$ mutations. Then the event $\{T > t\}$ occurs when at least one such event holds. Therefore, $\{T > t\} = \bigcup_{i=1}^{m} A_i(t)$. Because these events are not disjoint, since multiple edges may remain unmutated simultaneously, we calculate the probability of this union with the inclusion-exclusion principle:

$$\Pr(T > t) = \sum_{\emptyset \neq S \subseteq [m]} (-1)^{|S|+1} \Pr\Big(\bigcap_{i \in S} A_i(t)\Big)$$

The term with $|S| = 1$ adds the probability that a single edge remains unmutated, the terms with $|S| = 2$ subtract the probability that two edges remain unmutated simultaneously (since these were double-counted), and so on. Thus the alternating sum from the inclusion-exclusion principle exactly corrects for all overlaps among the events $A_i(t)$.

To evaluate $\Pr(\bigcap_{i \in S} A_i(t))$, let's consider one mutation event. Each mutation independently targets edge $i$ with probability $p_i$, where $\sum_{i=1}^{m} p_i = 1$. Hence, for a single mutation, the probability that it *does not* hit any edge in $S$ is

$$1 - \sum_{i \in S} p_i,$$

since the total probability of hitting any edge in $S$ is $\sum_{i \in S} p_i$. Because the $t$ mutations are independent, the probability that all $t$ of them avoid every edge in $S$ is obtained by multiplying this probability across all $t$ draws:

$$\Pr\Big(\bigcap_{i \in S} A_i(t)\Big) = \big(1 - \sum_{i \in S} p_i\big)^t$$

Intuitively, this is the probability that none of the $t$ mutations falls on any of the edges in $S$, i.e., that all edges in $S$ remain unmutated after $t$ mutation events. Applying this to Equation (S2), we get

$$\begin{aligned}
\mathbb{E}[T] &= \sum_{t=0}^{\infty} \Pr(T > t) \\
&= \sum_{\emptyset \neq S \subseteq [m]} (-1)^{|S|+1} \sum_{t=0}^{\infty} \Big(1 - \sum_{i \in S} p_i\Big)^t \\
&= \sum_{\emptyset \neq S \subseteq [m]} (-1)^{|S|+1} \frac{1}{\sum_{i \in S} p_i} \qquad \text{(geometric series with ratio } 1 - \sum_{i \in S} p_i \in [0, 1))
\end{aligned}$$

Finally, using $\frac{1}{x} = \int_0^\infty e^{-xu} du$ for $x = \sum_{i \in S} p_i$, the inclusion-exclusion identity $\prod_{i=1}^{m}(1 - z_i) = \sum_{S \subseteq [m]} (-1)^{|S|} \prod_{i \in S} z_i$, and substituting $z_i = e^{-p_i u}$ gives:

$$\mathbb{E}[T] = \int_0^\infty \Big[1 - \prod_{i=1}^{m}(1 - e^{-p_i u})\Big] du,$$

which is Equation (4) in the main text. □

**Lemma S1 (Equal branch lengths).** *If $p_i = \frac{1}{m}$ for all $i \in [m]$, then*

$$\mathbb{E}[T] = \int_0^\infty \left[1 - (1 - e^{-u/m})^m\right] du = mH_m$$

*Proof.*

$$\begin{aligned}
\mathbb{E}[T] &= \int_0^\infty \left[1 - (1 - e^{-u/m})^m\right] du && \text{(from Equation (S1))} \\
&= m \int_1^0 \frac{1 - (1 - x)^m}{x} (-dx) && (x = e^{-u/m}\text{, so } du = -\tfrac{m}{x} dx,\ u : 0 \to \infty \Rightarrow x : 1 \to 0) \\
&= m \int_0^1 \frac{1 - (1 - x)^m}{x} dx && \text{(flipping limits to remove the minus)} \\
&= m \int_0^1 \sum_{k=0}^{m-1} (1 - x)^k dx = m \sum_{k=0}^{m-1} \frac{1}{k+1} = mH_m && \text{(geometric series; integrating } (1-x)^k \text{ on } [0,1]\text{).}
\end{aligned}$$

**Lemma S2 (Expected total branch length of the Kingman coalescent trees).** *Let $L_{tot}$ be the total tree length (sum of all branch lengths) of the Kingman coalescent tree with n leaves and rate $\binom{k}{2}$ when there are k lineages. Then*

$$\mathbb{E}[L_{tot}] = 2 \sum_{j=1}^{n-1} \frac{1}{j} = 2\, H_{n-1}$$

*Proof.* Let $T_k$ be the waiting time until next coalescence, when $k$ lineages are present. The total tree length is the sum of the branch lengths contributed by each coalescent event:

$$L_{\text{tot}} = \sum_{k=2}^{n} k\, T_k$$

Under the coalescent model, $T_k$ follows an exponential distribution with rate $\binom{k}{2}$, hence $\mathbb{E}[T_k] = 1/\binom{k}{2}$ Taking expectations gives:

$$\mathbb{E}[L_{\text{tot}}] = \sum_{k=2}^{n} k\, \mathbb{E}[T_k] = \sum_{k=2}^{n} \frac{k}{\binom{k}{2}} = \sum_{k=2}^{n} \frac{2}{k-1} = 2 \sum_{j=1}^{n-1} \frac{1}{j} = 2\, H_{n-1}$$

□

#### S2 Supplementary Results

##### S2.1 Evaluation Criteria

*Pairwise Ancestral Accuracy.* Given the ground-truth tree $\mathcal{T}$ and an inferred tree $\mathcal{T}'$, we consider every pair of mutations $(a, b)$ that appear in both trees. For each tree, we determine whether mutation $a$ is ancestral to $b$, $b$ is ancestral to $a$, or if the two mutations are independent (i.e., neither is ancestral to the other). The pairwise ancestral relationship accuracy is then defined as the fraction of mutation pairs for which the inferred tree $\mathcal{T}'$ assigns the same ancestral relationship as the ground-truth tree $\mathcal{T}$. Formally,

$$\text{Accuracy} = \frac{|\{(a, b) : r_{\mathcal{T}}(a, b) = r_{\mathcal{T}'}(a, b)\}|}{\binom{M}{2}},$$

where $M$ is the number of mutations shared between $\mathcal{T}$ and $\mathcal{T}'$, and corresponds to the subset of mutations from $\mathcal{T}$ after subsampling. $r_{\mathcal{T}}(a, b) \in$ {ancestor, descendant, independent} denotes the ancestral relationship between $a$ and $b$ in tree $\mathcal{T}$.

##### S2.2 Perfect and 1-Dollo simulations

To evaluate the robustness of our bounds for perfect and 1-Dollo simulations, we first generated perfect phylogenies with $n \in \{2,3,4,5,6,10,15,20,25,30,40,50\}$ leaves. This corresponds to $(2n-1) \in \{3,5,7,9,11,19,29,39,49,59,79,99\}$ total nodes or clones in the tree. Since a full binary tree with $n$ leaves has $m = 2n-2$ edges, we generated $mH_m$ mutations and mapped each of these $mH_m$ mutations to a uniformly chosen edge in the tree. After assigning all mutations, we counted how many branches of the perfect phylogeny remained without any mutation. We simulated 10 replicates for each value of $n$. Figure S1 shows the distribution of the number of unmapped branches. The results indicate that using the perfect-phylogeny bound almost always results in every branch receiving at least one mutation.

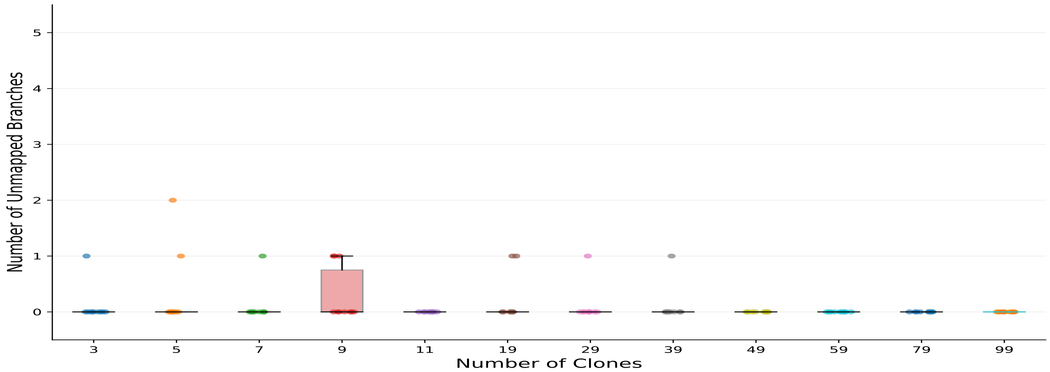


Fig. S1: Results on the number of unmapped branches for perfect phylogenies with $\{3,5,7,9,11,19,29,39,49,59,79,99\}$ clones.

##### S2.3 K-dollo phylogeny reconstruction accuracies for different loss probabilities

We simulated $k$-Dollo phylogenies with loss probabilities $p \in \{0.2, 0.3, 0.4\}$, following the procedure described in Section 3.1. Simulations were conducted for clone counts $\{3,5,7,9,11,19,29\}$, with the number of sampled mutations set as a multiple $x \in \{0.05, 0.1, 0.5, 0.7, 1, 2, 3\}$ of the perfect-phylogeny bound. For each combination of parameters, we generated 10 independent instances and used SPhyR [4] to reconstruct the corresponding $k$-Dollo phylogenies. Figures S2, S3, and S4 correspond to loss probabilities $p = 0.2$, $p = 0.3$, and $p = 0.4$, respectively, and show trends consistent with the $p = 0.1$ case described in the main text. However, as both the loss probability and $k$ increase, SPhyR [4] requires a larger number of sampled mutations to accurately reconstruct the phylogenies.

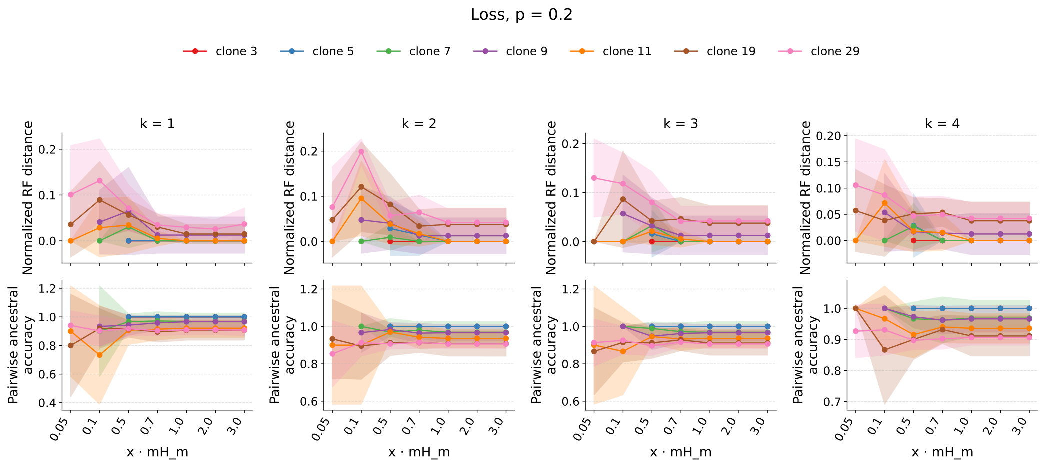


Fig. S2: Results for k-Dollo phylogeny reconstruction with SPhyR. The figure shows normalized phylogeny inference (top) and pairwise ancestral accuracy (bottom) across a range of $k$, clone numbers, and sampling rates for loss probability, $p = 0.2$.

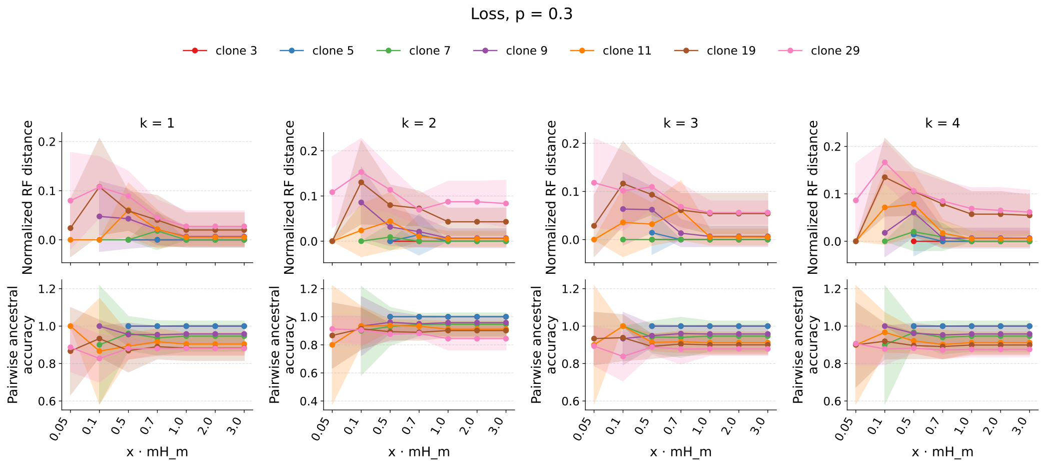


Fig. S3: Results for k-Dollo phylogeny reconstruction with SPhyR. The figure shows normalized phylogeny inference (top) and pairwise ancestral accuracy (bottom) across a range of $k$, clone numbers, and sampling rates for loss probability, $p = 0.3$.

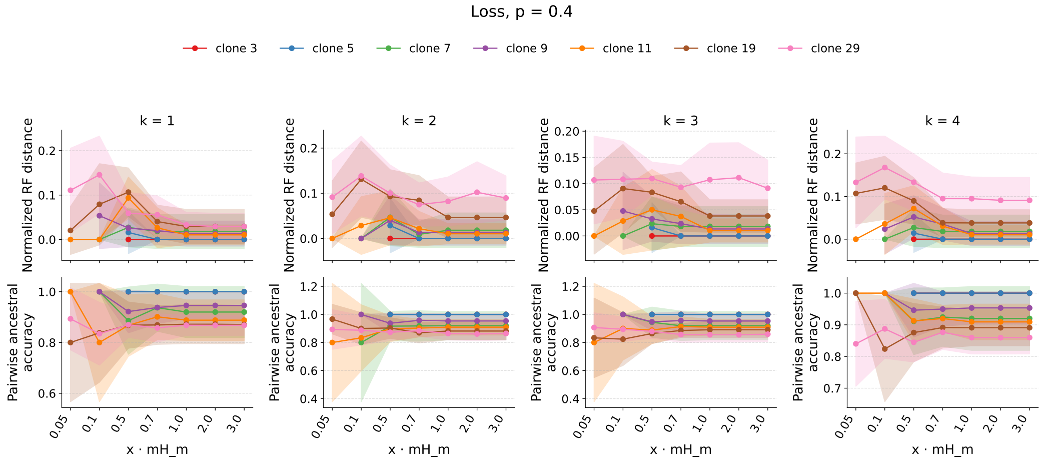


Fig. S4: Results for k-Dollo phylogeny reconstruction with SPhyR. The figure shows normalized phylogeny inference (top) and pairwise ancestral accuracy (bottom) across a range of $k$, clone numbers, and sampling rates for loss probability, $p = 0.4$.
